## Supplementary material for "Molecular quantitative trait loci in reproductive tissues impact male fertility in cattle": Supplement_NatComms_final.docx

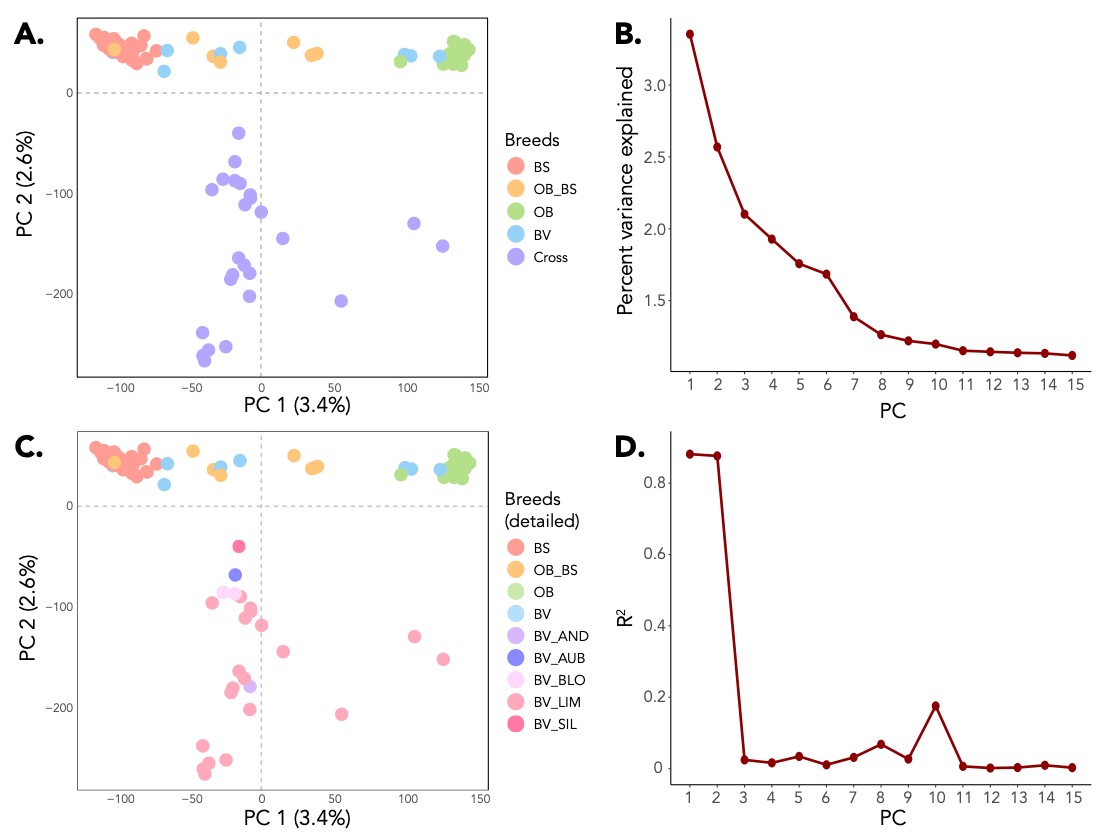

**Figure S1. Ancestry of 118 bulls from the molQTL cohort.** A.) PCA of LD-pruned WGS variants (366,090 variants total), with colors corresponding to the breed reported in the breed herdbook (BS = Brown Swiss; OB_BS = cross between Original Braunvieh and Brown Swiss; OB = Original Braunvieh; BV = Braunvieh) or “Cross”. B.) The percent variance explained for the first 15 PCs of the genotype PCA. C.) PCA of WGS variants, with colors corresponding to the specific breed reported. Abbreviations of the cross breeds are the following: AND = other (unknown); AUB = Aubrac; BLO = Blonde d'Aquitaine; LIM = Limousin; SIL = Silian (Simmental, Limousin and Angus). D.) Correlation between the PCs of the genotype PCA and the breed reported.

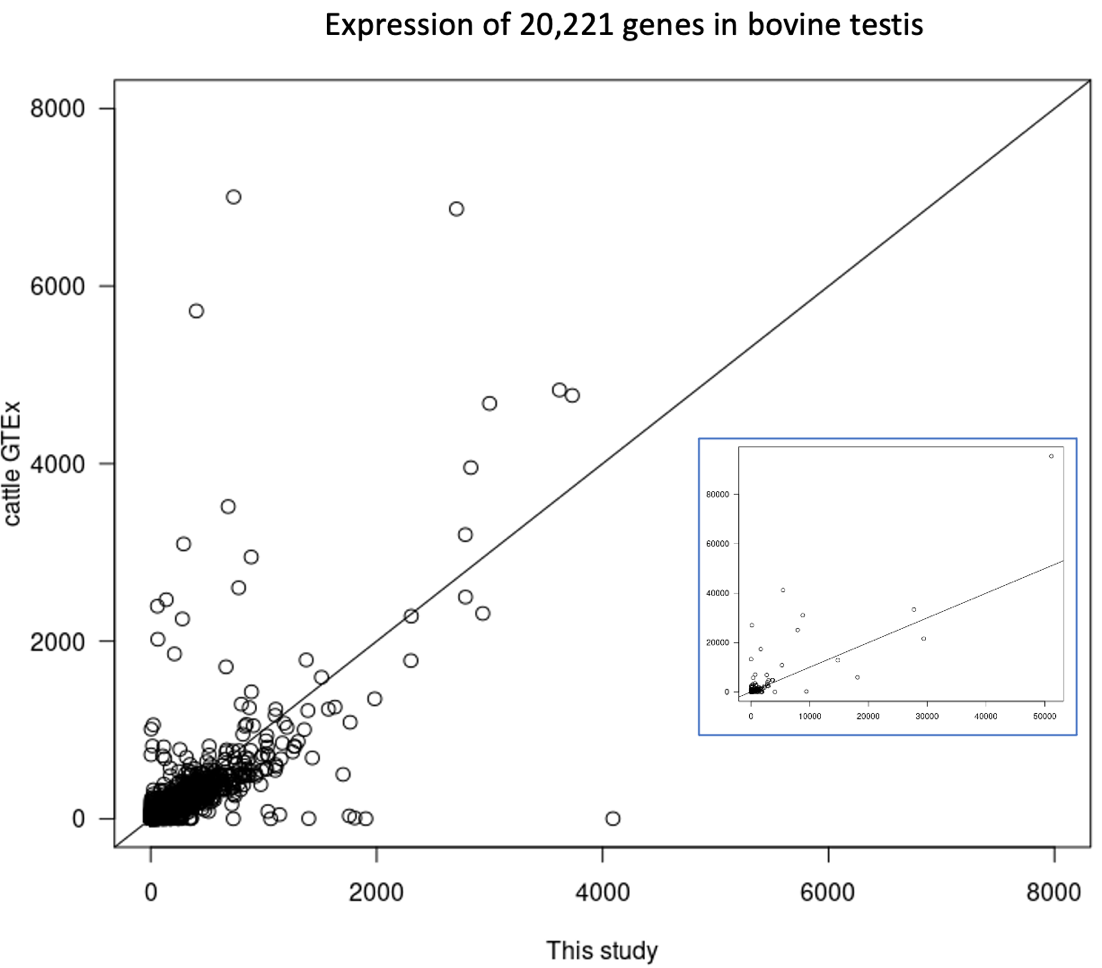

**Figure S2. Comparison of testis gene expression between this study and cattle GTEx.** Scatterplot of TPM values for 20,221 genes that are part of the current study and cGTEx. The axes are truncated at 8000. The solid line is a line through the origin. The inset shows the same plot without truncated axes. TPM estimates from cGTEx were obtained from the version 1.2 dataset deposited at zenodo (https://zenodo.org/record/7560235). Samples were filtered based on meta data to keep only post-pubertal taurine bulls. The following samples were retained: SRS3309609, SRS3309607, SRS3309608, SRS3021383, CRS013209, CRS013202, CRS013216, CRS013223, SRS734278, SRS485384.

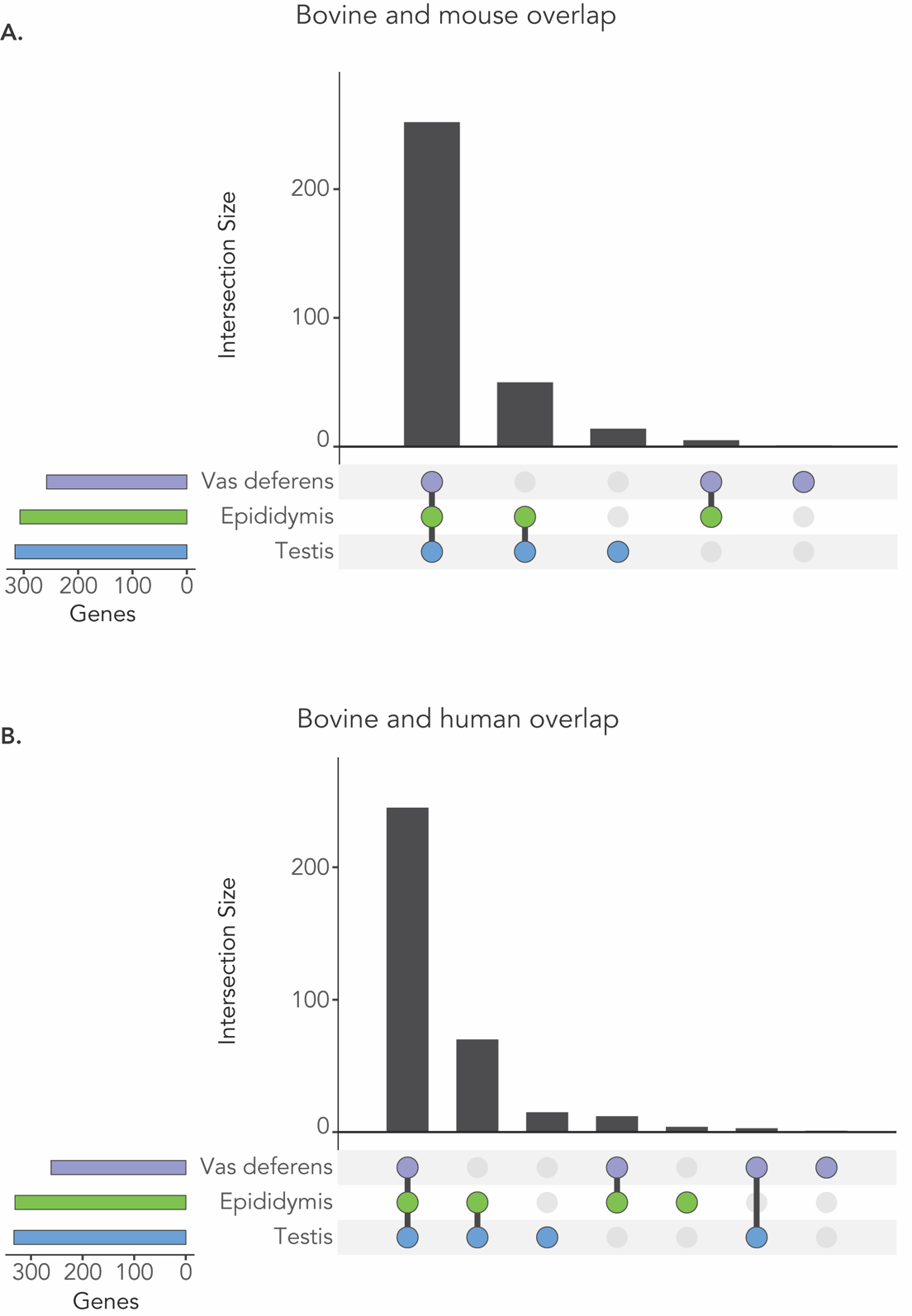

**Figure S3. Overlap of genes expressed in three bovine reproductive tissues with male reproductive tract-specific expressed genes in humans and mice.** We were able to unambiguously identify bovine orthologs for 380 of 720 reproductive tract-specific expressed genes in humans, and for 329 of 699 reproductive tract-specific expressed genes in mice [Robertson et al. (2019), https://doi.org/10.1186/s12915-020-00826-z]. A) UpsetR plot showing the expression pattern of the bovine orthologs of reproductive tract-specific expressed genes in mice. B) UpsetR plot showing the expression pattern of the bovine orthologs of reproductive tract-specific expressed genes in human.

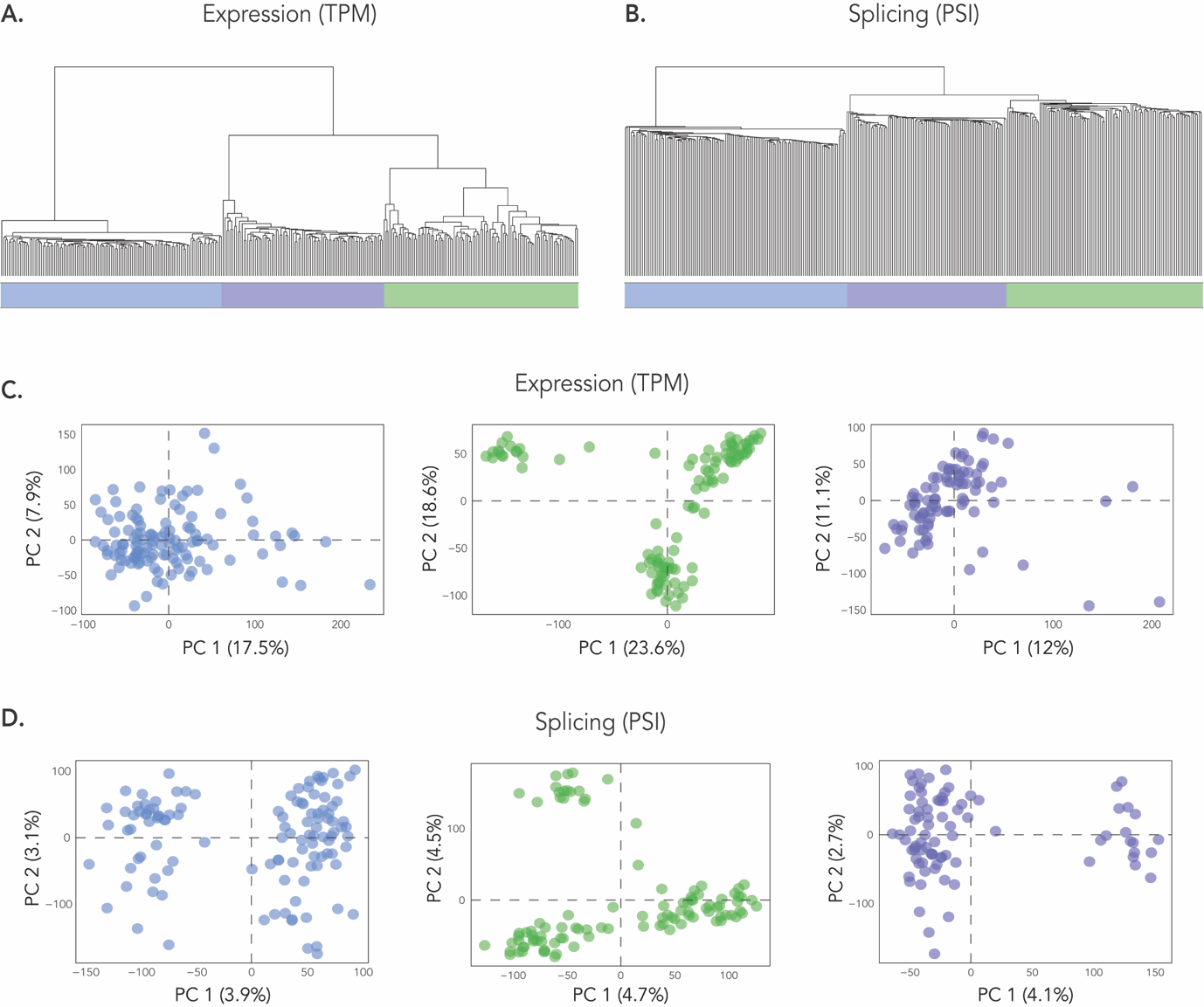

**Figure S4. Expression and splicing across male reproductive tissues.** A.) Hierarchical clustering from the r package *dendextend* for the normalized expression values (TPM) across the three tissues. Colored bars correspond to a sample’s tissue type (testis: blue; epididymis: green; vas deferens: purple). B.) Hierarchical clustering for the normalized splicing values (PSI) across the three tissues. Colored bars correspond to a sample’s tissue type. C.) Principal components analysis (PCA) of TPM values within individual tissues. The left plot is testis samples (blue), the middle plot is epididymis samples (green), and the right plot is vas deferens samples (purple). D.) PCA of PSI values within individual tissues. The left plot is testis samples (blue), the middle plot is epididymis samples (green), and the right plot is vas deferens samples (purple).

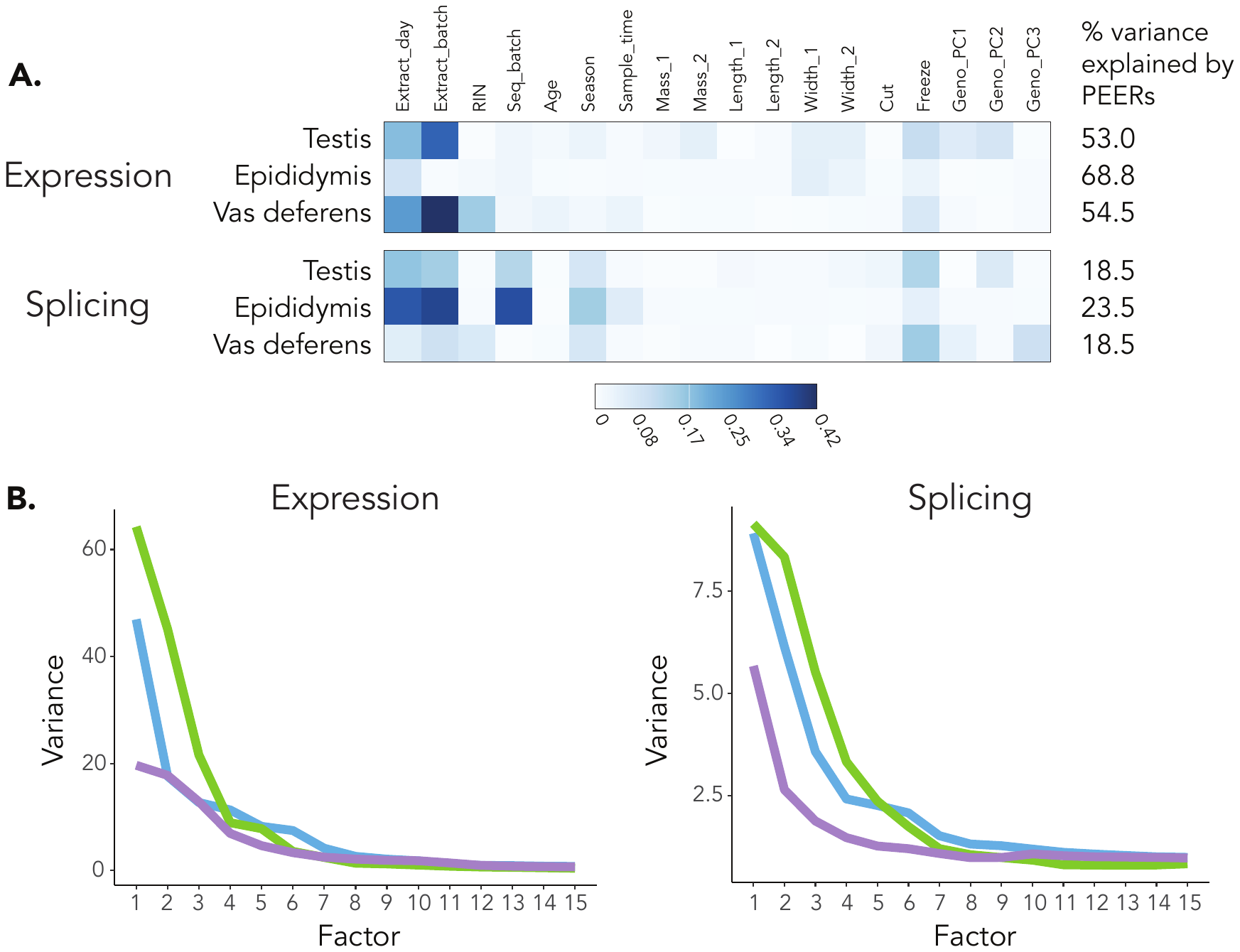

**Figure S5. PEER factors for expression and splicing phenotypes across tissues.** A.) Correlation between PEER factors and covariates for expression and splicing phenotypes across all tissues. The percent of variance in expression explained by the PEER factors for expression and splicing phenotypes for all tissues. B.) Posterior variance of the PEER factor weights across the tissues for expression and splicing values.

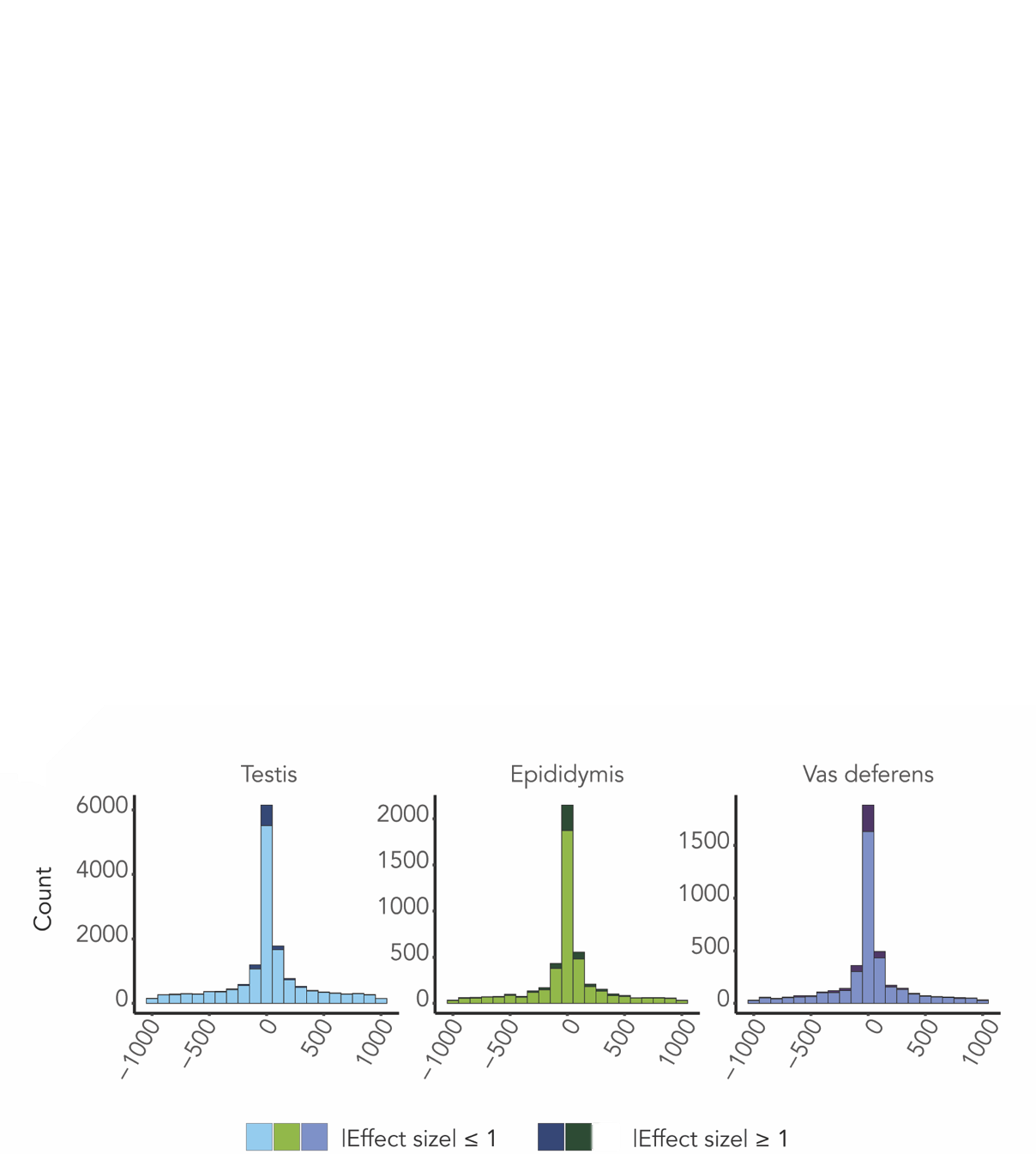

**Figure S6. eQTL and sQTL characteristics across the three tissues.** Distribution of eQTL from the TSS for all three reproductive tissues. Darker colors represent large effect eQTL (|slope_aFC >= 1|), while lighter colors represent small effect eQTL (|slope_aFC <= 1|).

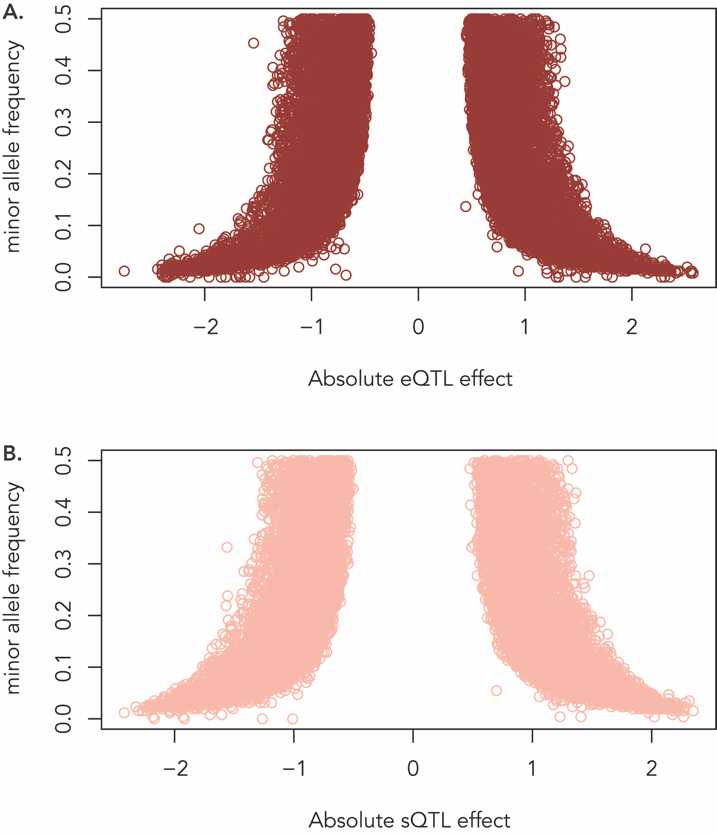

**Figure S7. Minor allele frequency and effect size of eQTL and sQTL.** A.) Minor allele frequency and effect size of eQTL identified across the three tissues. B.) Minor allele frequency and effect size of sQTL identified across the three tissues.

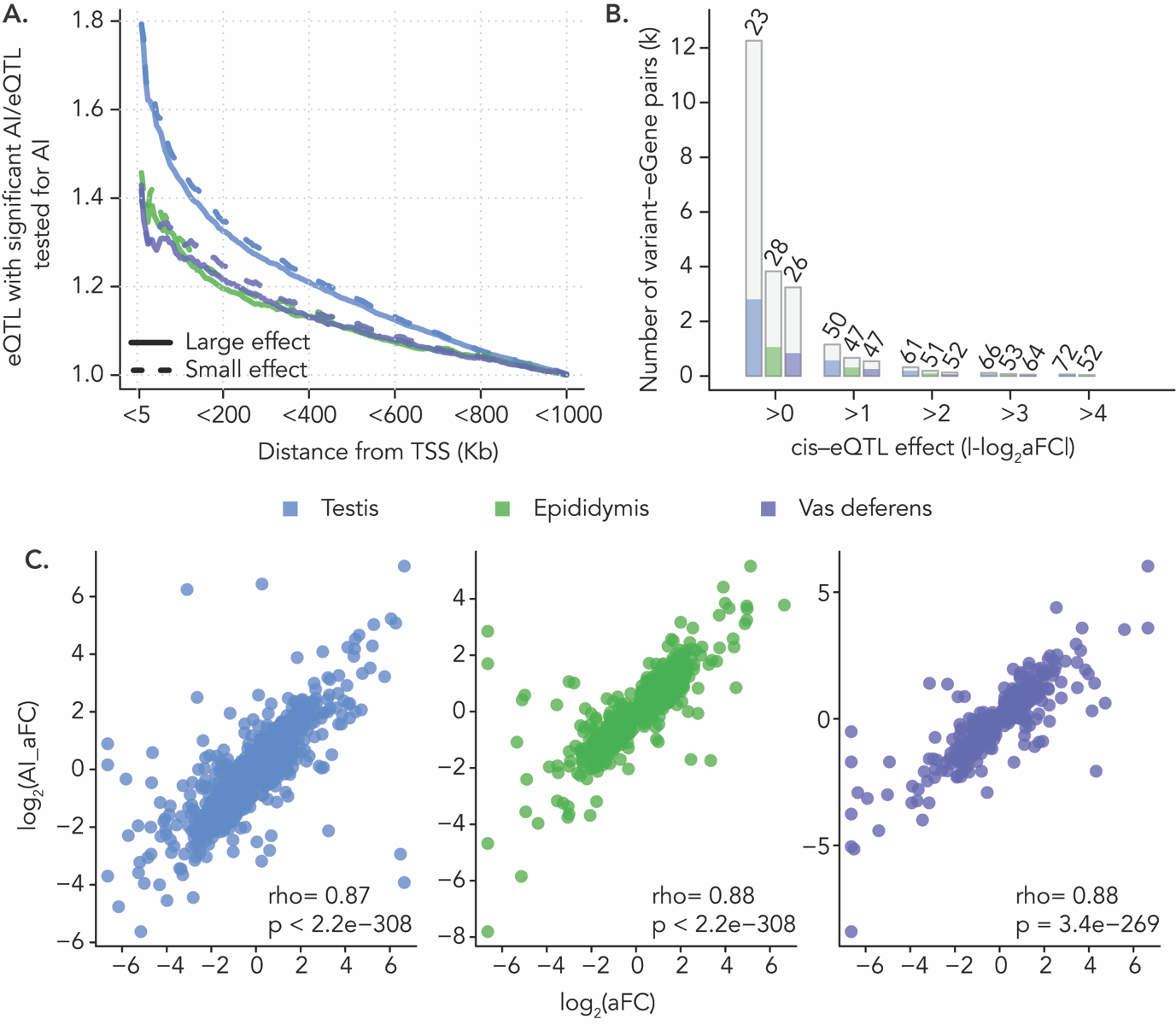
**Figure S8. Analysis of Allelic imbalance in expression of eGenes to assess cis-regulatory activity of linked eQTL.** A.) Enrichment of cis-eQTL associated with significant Allelic imbalance grouped by their proximity to TSS (in bins of 5kb) in the three tissues compared to random expectation. Data for large effect (|log_2_ slope_aFC|>=1) and small effect (|log_2_ slope_aFC| < 1) eQTL are presented separately. B.) Number (in thousands) of variant-gene pairs tested for allelic imbalance (height of the bar) and with significant allelic imbalance (height of colored bar) grouped by their effect sizes (|log_2_ slope_FC|). The percentage of cis-eQTL with significant allelic imbalance are indicated at the top of the bar. C.) Correlation (Spearman’s rho) between the cis-eQTL effect estimated as slope_aFC and effect estimated from the allelic imbalance data (AI_aFC). Only variant-gene pairs showing significant allelic imbalance (FDR <0.05) were considered. Spearman’s correlations and their significances are indicated at the bottom.

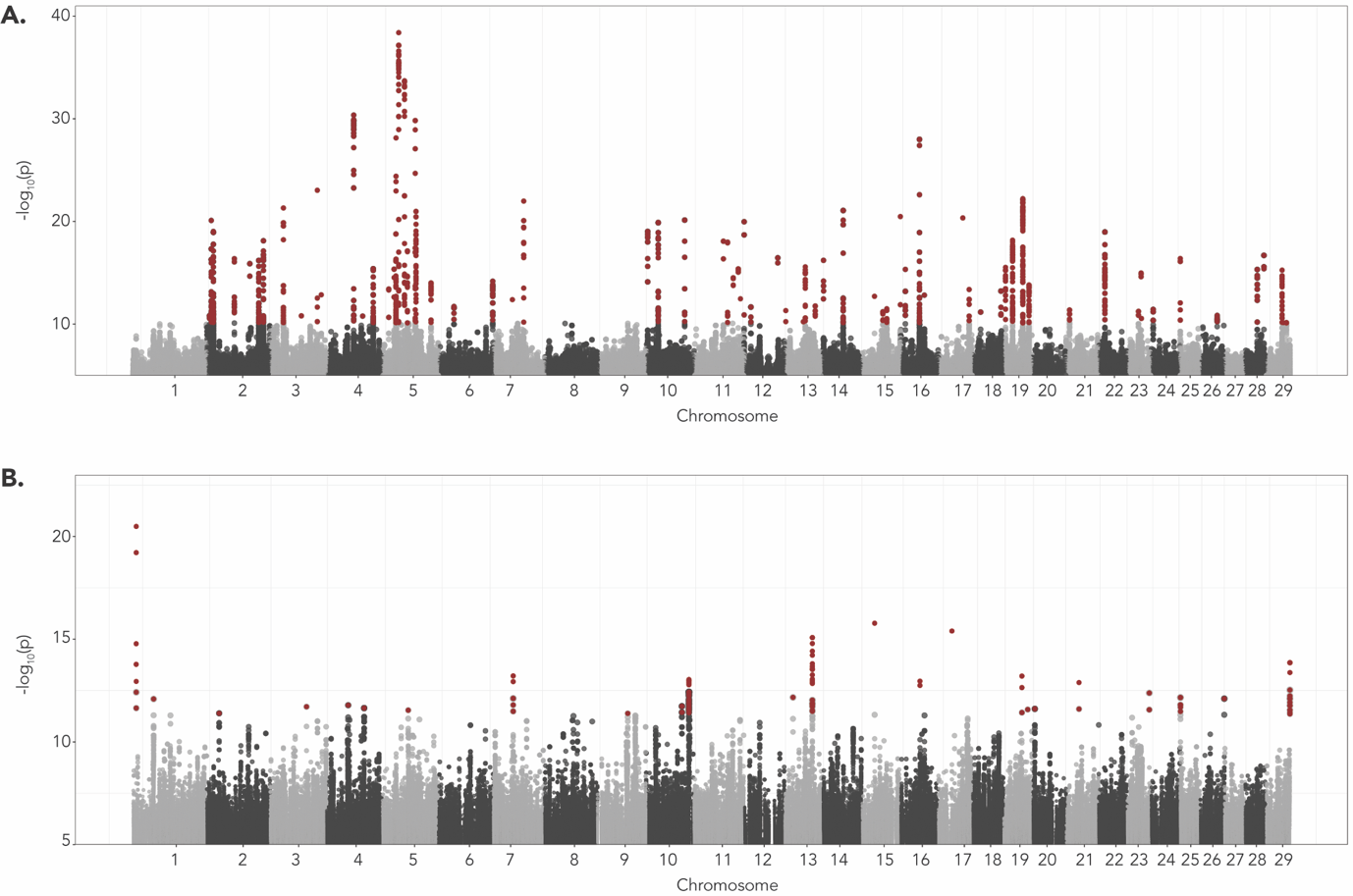

**Figure S9. Trans molQTL mapping in testis tissue.** A) Manhattan plot representing the results of trans-eQTL mapping. B) Manhattan plot representing the results of trans-sQTL mapping. Red color indicates significant e/sVariants.

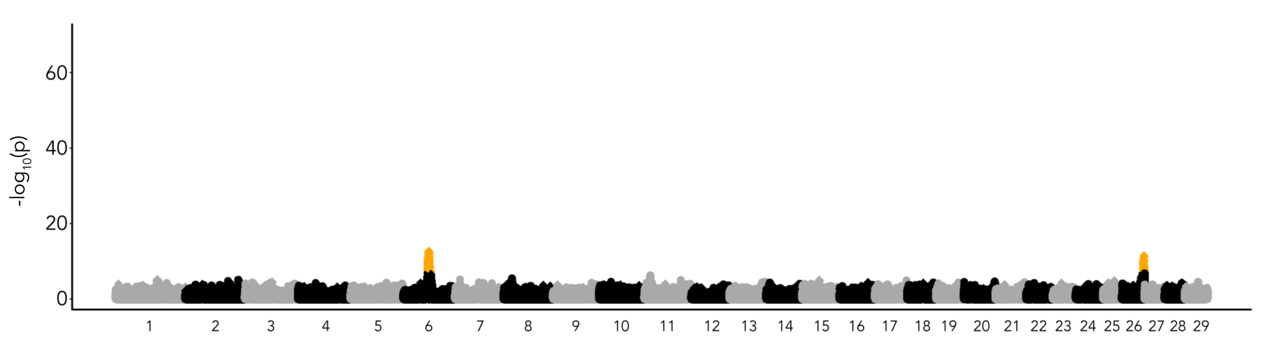

**Figure S10. Additive GWAS of male fertility.** Manhattan plot showing genome-wide association between male fertility and imputed sequence variants from additive association models. Orange dots indicate variants that are significantly associated with male fertility at a significance threshold of 5e-08.

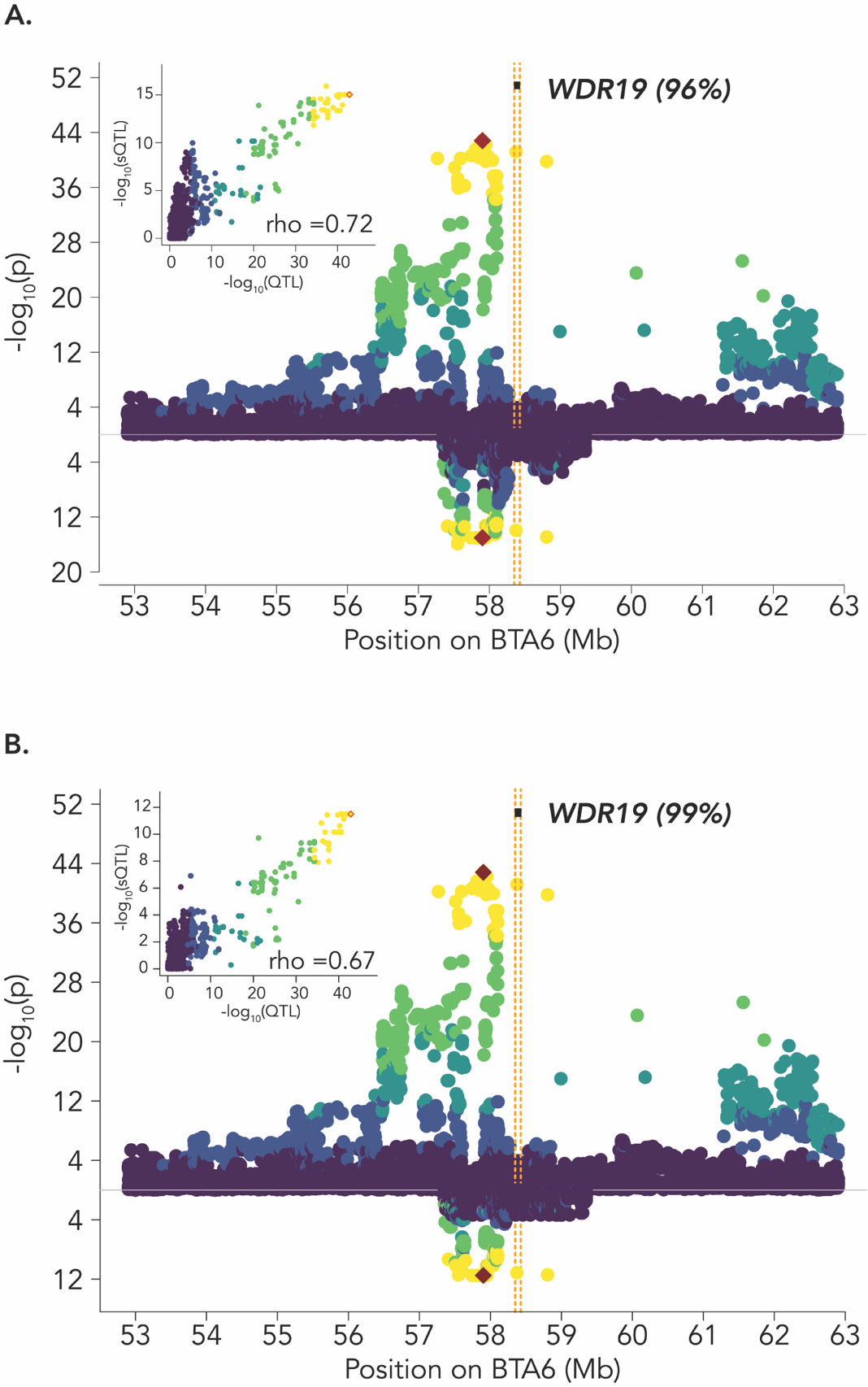

**Figure S11. Molecular *WDR19* phenotype overlap with a male fertility QTL.** A.) Mirrored plots of –log_10_(p)-values from association testing between imputed sequence variants and bull fertility (top) and splicing phenotypes for intron cluster 63814 spanning the splice junction 58,373,894-58,374,821 of the gene *WDR19* (bottom) in epididymis (A) and intron cluster spanning the same splice junction of the gene WDR19 in Vas deferens (B). The inset plots show the correlation between GWAS and sQTL –log_10_(p)-values. Color indicates the linkage disequilibrium (R^2^) between the most likely colocalized variant in the two tissues (6:57900948_G_A, red colored) and all other variants.

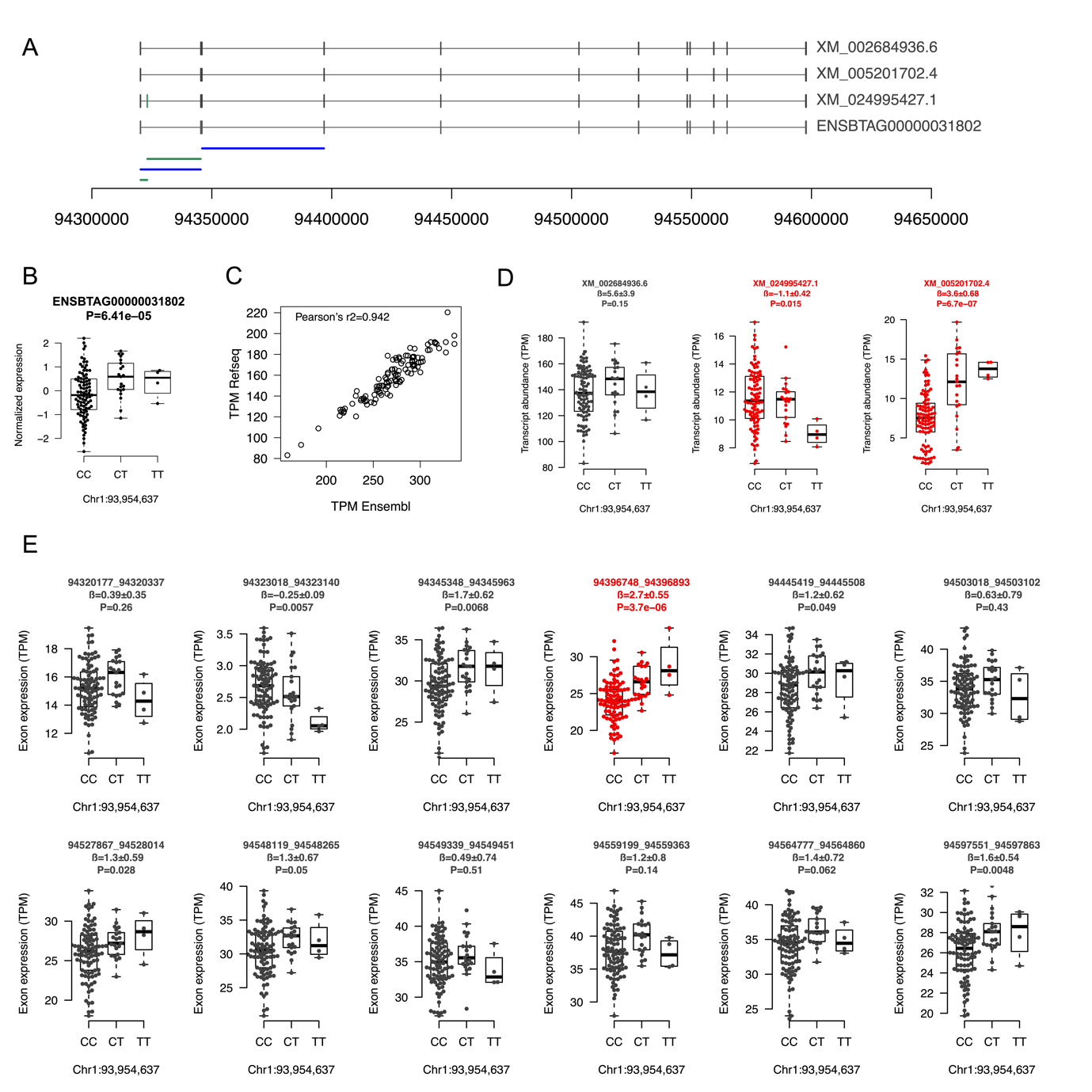

**Figure S12. Alternative splicing and expression of SPATA16.** A) Comparison of the structure between three SPATA16 isoforms (XM_002684936.6, XM_005201702.4, XM_024995427.1) that are annotated in Refseq (version 106) and the canonical SPATA16 transcript (ENSBTAG00000031802) annotated from Ensembl (version 104). Boxes represent exons. The exon private to XM_024995427.1 is coloured in green. Blue and green lines below the gene structures indicate the four splicing junctions of the significant intron cluster. Green colour indicates two junctions that involve the exon private to XM_024995427.1. B) Effect of the Chr1:93954637 variant on the expression of ENSBTAG00000031802. The P value is from a linear regression of TPM values on the genotype (coded as 0, 1, 2) while accounting for age, RIN, 10 PEER factors and three principal components of a genomic relationship matrix. C) The gene-level TPM values of SPATA16 are highly correlated (Pearson's rho=0.942) between the Ensembl and Refseq annotation. D) Effect of the Chr1:93954637 variant on transcript-level expression estimates. Transcripts are from the Refseq annotation. Red colour indicates significant transcripts. E) Effect of the Chr1:93954637 variant on exon-level expression estimates. Exons are from the Refseq annotation. Red colour indicates significant exons. The P values presented in D) and E) are from a linear regression of TPM values on the genotype (coded as 0, 1, 2) while accounting for age, RIN and three principal components of a genomic relationship matrix.

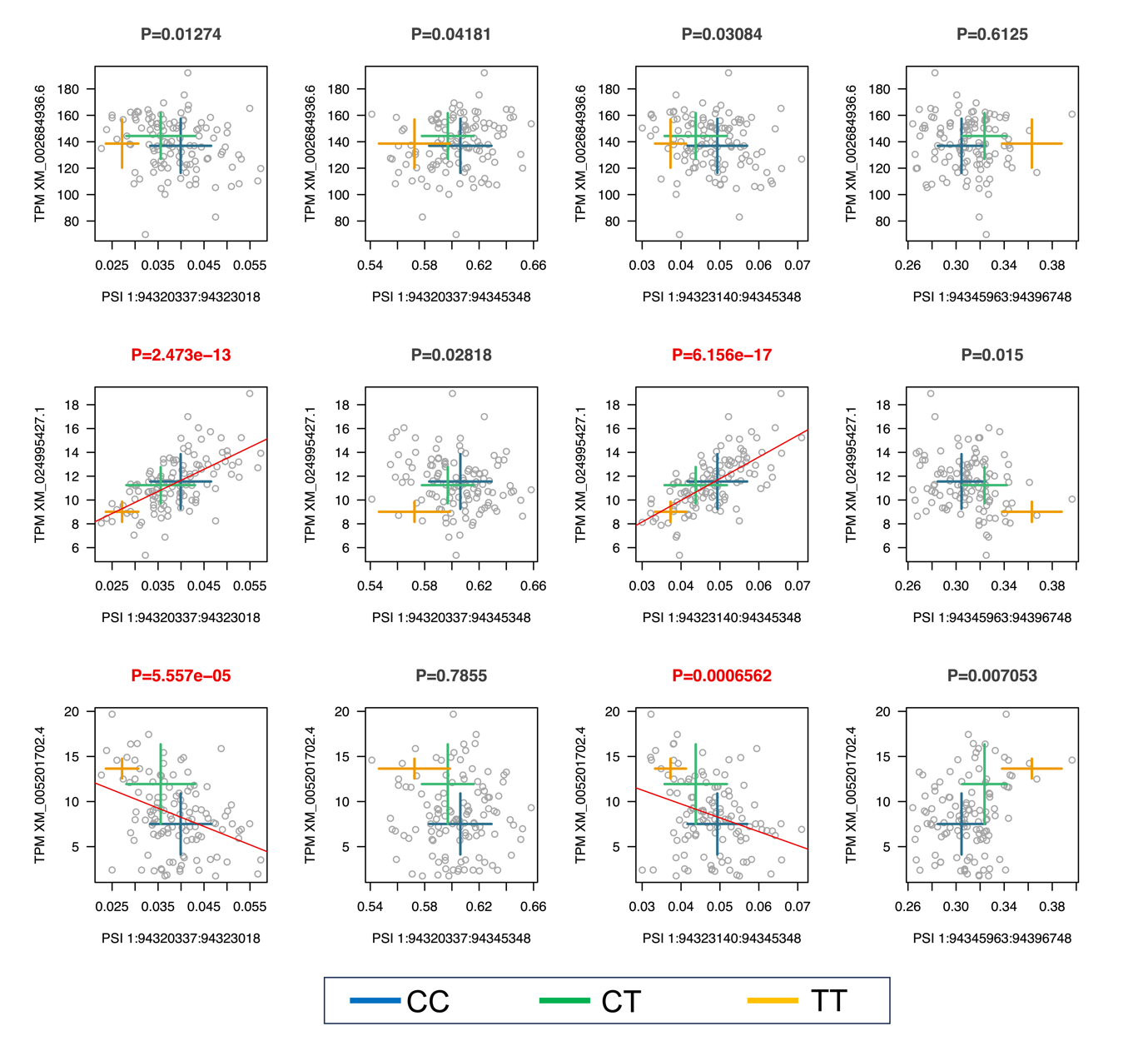
**Figure S13. SPATA16 TPM and PSI correlations.** Correlation between the expression (quantified in TPM) of the three SPATA16 isoforms (XM_002684936.6, XM_005201702.4, XM_024995427.1) that are annotated in Refseq and the percent-spliced-in (PSI)-values of the four splice junctions of the significant intron cluster. P values were from a linear regression of TPM on PSI. Red colour indicates significant transcript-splice junction relationships and the red line is the corresponding regression line. The two splice junctions that are significanly associated with the expression of XM_005201702.4 and XM_024995427.1 involve the exon private to XM_024995427.1. Different colours indicate mean ± standard deviations for PSI and TPM values for different Chr1:93954637 genotypes.

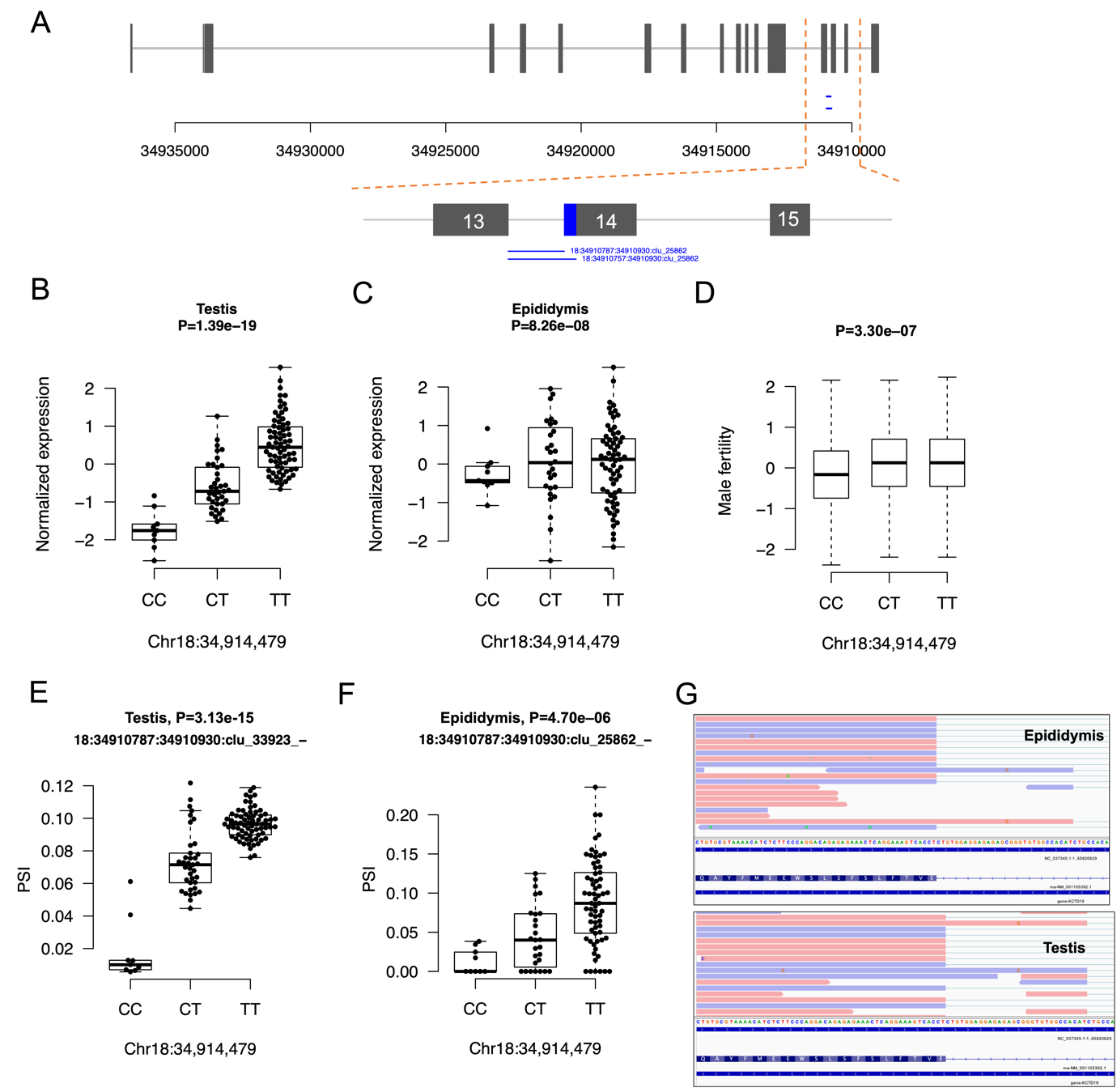
 **Figure S14. Alternative splicing and expression of KCTD19.** A) Structure of KCTD19. Boxes represent exons. Blue lines indicate two splice junctions within a significant intron cluster. The utilization of splice junction 34910787:34910930 adds 30 basepair coding sequence to the 14th exon (indicated with the blue box preceding exon 14). B) and C) Effect of the Chr18:34914479 variant on the expression of KCTD19 in testis and epididymis. P values are from the nominal eQTL mapping. D) Effect of the Chr18:34914479 variant on male fertility. The P value is from the non-additive GWAS model. E) and F) Effect of the Chr18:34914479 variant on KCTD19 splicing in testis and epididymis. The P values are from the nominal sQTL mapping. G) Representative IGV screenshots of epididymis and testis RNA sequencing alignments validate the alternative start of exon 14.

**Table S2. Genotype variants considered after different filtering methods**. Variants remaining after different filtering methods. MAF filters were applied separately for each subset of samples.

| **# Variants after filtering** | **Number of samples (total)** | **No filtering** | **HWE and missingness** | **Imputation accuracy > 0.5** |
| --- | --- | --- | --- | --- |
|  | 118 | 29,660,795 | 29,238,235 | 21,501,032 |
| **MAF**  **filters** | **Tissue** | **MAF ≥ 0.5%** | **MAF ≥ 1.0%** | **MAF ≥ 5.0%** |
|  | Testis (117) | 19,964,363 | 18,919,356 | 14,192,337 |
|  | Epididymis (103) | 19,704,122 | 18,606,598 | 14,041,832 |
|  | Vas deferens (84) | 21,041,956 | 19,367,827 | 14,208,661 |

**Table S3. RNA read alignment.** Information on the RNA sequence data for all tissue. The average and the standard deviation are reported for each measure.

| Tissue | RIN | # Aligned RNA reads | # Aligned RNA reads (WASP) |
| --- | --- | --- | --- |
| Testis | 9.0 ± 0.5 | 257,050,199 ± 35,278,398 | 240,544,139 ± 32,870,352 |
| Epididymis | 8.4 ± 0.9 | 283,746,666 ± 35,375,776 | 267,645,035 ± 32,952,821 |
| Vas deferens | 5.9 ± 1.1 | 262,089,072 ± 25,546,308 | 251,031,216 ± 24,272,517 |

**Table S4. Types of expressed and spliced genes, and types of eGenes and sGenes**. Breakdown of the number and proportion of expressed and spliced genes (after filtering), and eGenes and sGenes (after molQTL discovery), by type per tissue. “Other” includes pseudogenes and various small RNAs.

|  | **Type** | **Testis** | **Epididymis** | **Vas deferens** |
| --- | --- | --- | --- | --- |
| **Expressed genes** | Protein coding | 18,149 (89.7%) | 17,815 (87.4%) | 16,809 (88.2%) |
|  | Long non-coding | 933 (4.6%) | 1,002 (4.9%) | 831 (4.4%) |
|  | Other | 1,140 (5.7%) | 1,559 (7.7%) | 1,423 (7.4%) |
| **Spliced genes** | Protein coding | 14,087 (95.7%) | 13,470 (96.0%) | 12,364 (96.7%) |
|  | Long non-coding | 603 (4.1%) | 530 (3.8%) | 389 (3.0%) |
|  | Other | 34 (0.2%) | 26 (0.2%) | 27 (0.3%) |
| **eQTL** | Protein coding | 10,376 (93.0%) | 3,974 (91.3%) | 3,630 (93.3%) |
|  | Long non-coding | 497 (4.5%) | 202 (4.7%) | 159 (4.1%) |
|  | Other | 291 (2.5%) | 171 (4.0%) | 100 (2.6%) |
| **sQTL** | Protein coding | 6,735 (96.2%) | 2,571 (96.7%) | 1,660 (96.7%) |
|  | Long non-coding | 247 (3.5%) | 84 (3.1%) | 51 (3.0%) |
|  | Other | 18 (0.3%) | 7 (0.2%) | 7 (0.3%) |

**Table S5. Expression of differentially spliced genes.** Median gene expression (TPM) for differentially spliced genes

|  | Testis | Epididymis | Vas deferens |
| --- | --- | --- | --- |
| 11,474 genes with alternative splicing in all tissues | 17.37 | 15.89 | 20.18 |
| Genes with no alternative splicing | 1.73 (5,523 genes) | 1.91 (6,355 genes) | 2.13 (6,271 genes) |
